# Altered cohesin dynamics and histone H3K9 modifications contribute to mitotic defects in the *cbf11Δ* lipid metabolism mutant

**DOI:** 10.1101/2022.10.17.512562

**Authors:** Akshay Vishwanatha, Jarmila Princová, Patrik Hohoš, Róbert Zach, Martin Převorovský

**Author notes:** Whitney 7, Department of Medicine, 1300 York Avenue, New York, NY 10065, USA.

## Abstract

Mitotic fidelity is crucial for the faithful distribution of genetic information into the daughter cells. Many fungal species, including the fission yeast *Schizosaccharomyces pombe*, undergo a closed form of mitosis, during which the nuclear envelope does not break down. In *S. pombe* numerous processes have been identified that contribute to successful completion of mitosis. Notably, perturbations of lipid metabolism can lead to catastrophic mitosis and the “cut” phenotype. It was suggested that these mitotic defects are caused by insufficient membrane phospholipid supply during the anaphase nuclear expansion. However, it is not clear whether additional factors are involved. In this study we characterized in detail the mitosis in an *S. pombe* mutant lacking the Cbf11 transcription factor, which regulates lipid metabolism genes. We show that in *cbf11Δ* cells mitotic defects appear already prior to anaphase, before the nuclear expansion begins. Moreover, we identify altered cohesin dynamics and centromeric chromatin structure as additional factors affecting mitotic fidelity in cells with disrupted lipid homeostasis, providing new insights into this fundamental biological process.

## INTRODUCTION

Mitosis is an elaborate process critical for cell reproduction, and the fidelity of mitosis is crucial for the faithful distribution of genetic information into the daughter cells. Mitotic defects can result in aneuploidy, as frequently seen in cancer cells, or in catastrophic mitosis and cell death. As such, mitosis is tightly controlled and regulated at many different levels (Bähler, 2005; Cullati and Gould, 2019; Grallert et al., 2015). The fission yeast *Schizosaccharomyces pombe* has been instrumental in identifying the factors and processes involved in mitosis, and in defining their effects on mitotic fidelity (Hayles and Nurse, 2018; Hayles et al., 2013). *S. pombe* features regional, repetitive, and heterochromatinized centromeres resembling the centromeres of human cells. The special chromatin structure at the centromeres is important for proper kinetochore establishment and its attachment to the mitotic spindle (Roy et al., 2013; Tong et al., 2019). In post-replication chromosomes, sister chromatids are held together by cohesin rings, which in fission yeast are removed predominantly at the onset of anaphase to allow chromosome segregation (Peters et al., 2008). Unlike the human cells, *S. pombe* has a fully closed mitosis, during which the nuclear membrane does not break down and the spindle forms and elongates within the confines of the nuclear envelope (Zhang and Oliferenko, 2013).

Intriguingly, chemical inhibitors or genetic manipulations perturbing lipid metabolism have also been shown to affect mitotic fidelity in *S. pombe*. Inhibition of the fatty acid synthase or downregulation of acetyl coenzyme A carboxylase result in a form of catastrophic mitosis called the “cut” (<u>c</u>ell <u>u</u>ntimely torn) phenotype. In this lethal event, the cell nucleus is bisected by the cytokinetic septum before mitosis is completed (Makarova et al., 2016; Saitoh et al., 1996; Takemoto et al., 2016; Zach and Převorovský, 2018). Recent studies have highlighted the importance of rapid nuclear envelope expansion during anaphase for successful mitosis. The surface area of the nucleus enlarges by ∼30% in anaphase, and this process requires an adequate supply of membrane phospholipids. Failure to expand the nucleus is associated with mitotic spindle bending and breaking, chromosome segregation defects, and/or the “cut” phenotype. Consequently, it has been postulated that insufficient membrane phospholipid supply is the key factor underlying the mitotic defects observed in cells with perturbed lipid metabolism (Makarova et al., 2016; Takemoto et al., 2016).

Cbf11 is a CSL family transcription factor that directly regulates several lipid metabolism genes in *S. pombe*. Cells lacking Cbf11 show defects in cell cycle progression, including the “cut” phenotype. The *cbf11Δ* cells also show decreased content of storage lipid droplets, which would be consistent with a decreased capacity for membrane phospholipid production and, therefore, being prone to catastrophic mitosis (Převorovský et al., 2015; Převorovský et al., 2016). However, the deletion of *cbf11* was shown to exert intriguing genetic interactions with mutations in cohesin loaders, a chromatin modification complex, and kinetochore-related factors (Chen et al., 2012; Guo et al., 2014; Rallis et al., 2017). These findings suggested that there may be additional factors contributing to the occurrence of mitotic defects in cells with perturbed lipid metabolism. In this study, we set out to characterize in detail the mitotic progression in *cbf11Δ* cells and to explore the previously identified genetic interactions mentioned above. We report altered cohesin occupancy and histone H3K9 modifications at the centromeres, accompanied by an increased propensity for chromosome loss. Furthermore, we identify cohesin dynamics and chromatin structure as factors likely contributing to mitotic fidelity in the *cbf11Δ* lipid metabolism mutant.

## RESULTS

### Cells lacking Cbf11 are prone to aberrant mitotic outcomes

To monitor the dynamics and outcomes of mitosis in the *cbf11Δ* mutant, we performed live-cell time-lapse microscopy using strains with fluorescently tagged histone H3 (Hht2-GFP) and alpha-tubulin (mCherry-Atb2) (Syrovatkina and Tran, 2015) to visualize the chromosomes and mitotic spindle, respectively. Using wild-type (WT) and *cbf11Δ* cells, we detected 3 different mitotic outcomes: 1) normal successful mitosis concluded by the formation of two daughter cells with equally segregated chromosomes, 2) catastrophic mitosis in the form of the “cut” phenotype with unequal chromosome distribution in the daughter cells, and 3) nuclear displacement which results in the formation of a viable diploid daughter cell and a non-viable anucleate daughter cell (**Fig. 1A, B**). During the 336 minutes of the experiment, 51% of WT cells underwent mitosis, with only ∼6% of these showing aberrant mitotic outcomes (n = 263 total cells). In contrast, during the same period, only 13% of *cbf11Δ* cells underwent mitosis, but ∼31% of those showed aberrant mitotic outcomes (n = 333 total cells) (**Fig. 1C**). The lower percentage of mitotic cells in the observed *cbf11Δ* population might be linked to the slow-growth phenotype of this mutant. Interestingly, the nuclear displacement could explain the previously reported continual emergence of non-sporulating diploid subpopulations in *cbf11Δ* cultures (Převorovský et al., 2009). A similar, actin-dependent, nuclear displacement and evasion of catastrophic mitosis coupled with diploidization was recently documented in several other fission yeast “cut” mutants (Yukawa et al., 2021).

**Figure 1.**
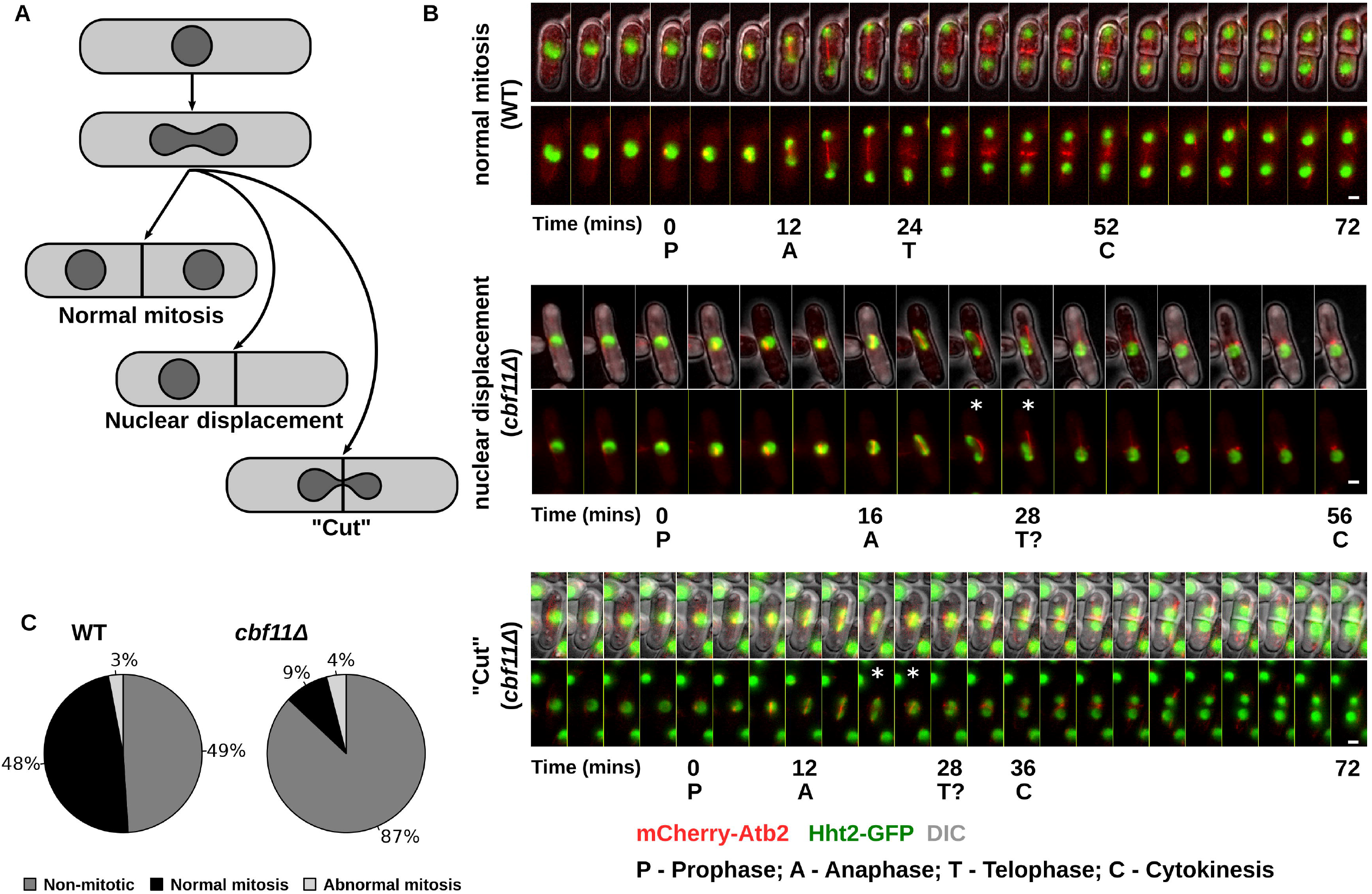
Mitotic outcomes in WT and *cbf11Δ* cells. **(A)** A diagram of possible mitotic outcomes in WT and *cbf11Δ* cells. **(B)** Representative examples of normal and aberrant mitotic timecourses. Bottom panels show chromosomes in green and microtubules in red; top panels show an overlay with DIC images to visualize cell boundaries and septa. The beginning of individual mitotic phases is indicated below the respective microscopic frames; asterisks indicate spindle defects. Bar represents 1 μm. **(C)** Quantification of mitotic outcomes during the course of the experiment.

### Cells lacking Cbf11 show aberrant mitotic timing and spindle dynamics

To better understand the nature of mitotic defects in *cbf11Δ* cells, we performed a more detailed analysis of our live-cell microscopy dataset from **Fig. 1**. We randomly selected 10 cells from the categories “WT normal”, “*cbf11Δ* normal” and “*cbf11Δ* abnormal”, and measured the timing and dynamics of nuclear division, chromosome segregation, spindle elongation, and cytokinesis (**Fig. 2** and **Fig. S1**). WT cells displayed rapid and relatively uniform mitotic progression, with spindle formation starting ∼12-18 minutes before the onset of anaphase (**Fig. 2A**) and median mitotic duration being 26 min (**Fig. 2E**). In contrast, the timing of mitotic progression was markedly perturbed in the *cbf11Δ* mutant. In these cells, the interval between spindle formation and anaphase onset was more variable and typically longer than in WT (∼10-30 min; **Fig. 2B**), suggesting possible problems with the initial attachment of chromosomes to spindle microtubules. Furthermore, in mutant cells that completed mitosis successfully, the spindle elongation was slower (**Fig. 2C, Fig. S1A**), and total mitotic duration was longer and more variable compared to WT (**Fig. 2E**).

**Figure 2.**
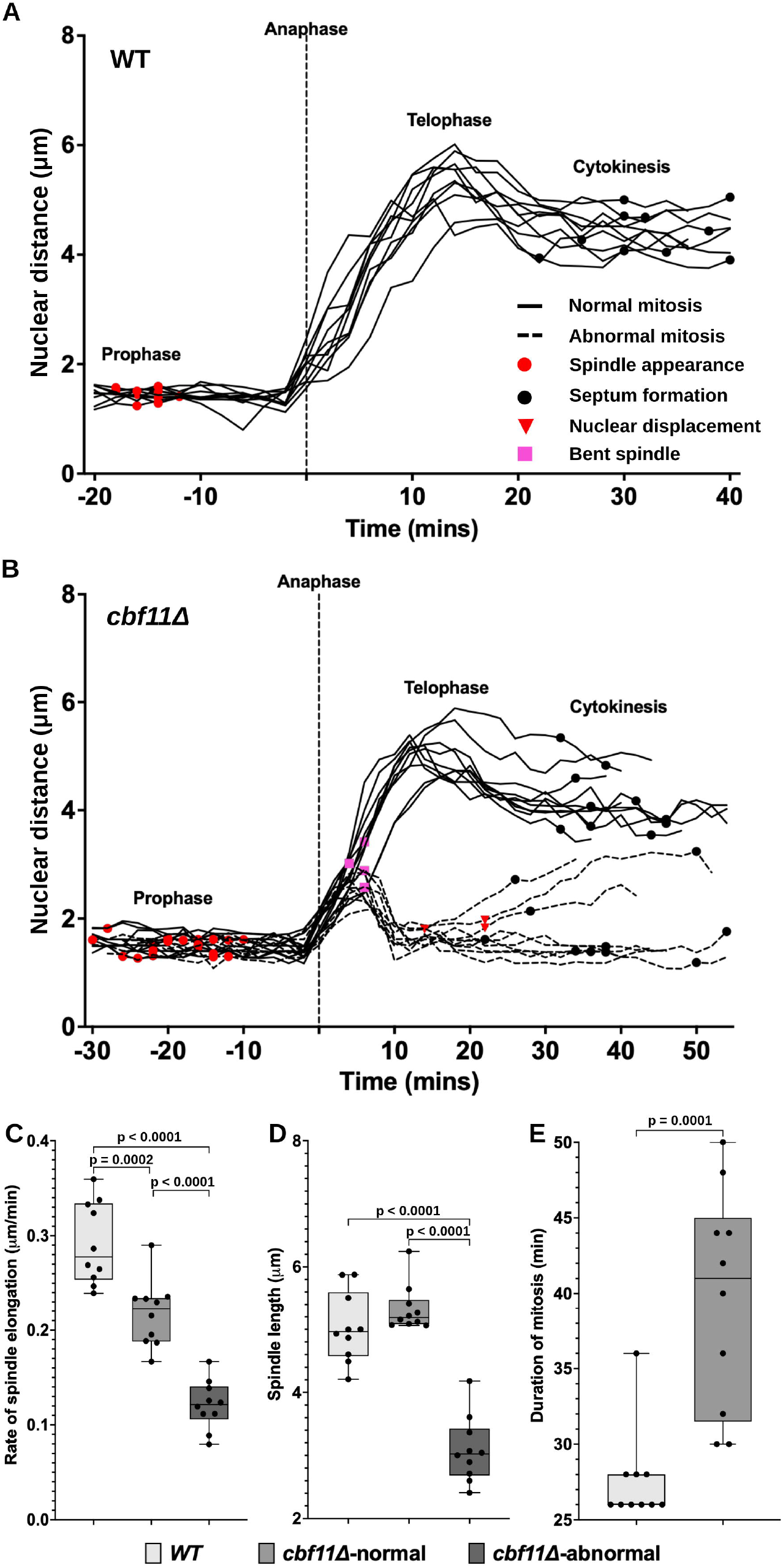
Dynamics of mitotic progression in WT and *cbf11Δ* cells. **(A, B)** Timing of mitotic events in WT and *cbf11Δ* cells, respectively. The distance between opposite nuclear edges during mitosis is shown, and important mitotic events are indicated. Ten cells were analyzed for each group (WT, *cbf11Δ* normal, *cbf11Δ* abnormal). The legend in panel (A) also applies to panel (B). **(C)** Rate of spindle elongation during anaphase. **(D)** Maximum spindle length during mitosis. **(E)** Total duration of successfully completed mitoses. The shared legend for panels (C-E) is shown at the bottom. Significance of differences between groups was determined by one-way ANOVA followed by the Bonferroni multiple comparison test (C, D), or by a two-tailed unpaired t-test (E).

In the case of *cbf11Δ* mitoses with aberrant outcomes, the spindle elongation rate was even more impaired (**Fig. 2C, Fig. S1B**), and the maximum spindle length reached before its disassembly was markedly shorter than in either WT or successful *cbf11Δ* mitoses (**Fig. 2D**). Notably, the nuclear displacement phenotype was associated with mid-anaphase spindle bending and/or detachment of one of the daughter chromosome masses from the spindle, and subsequent spindle disassembly and merger of the two daughter chromosome masses into one diploid nucleus (**Fig. 1B, Fig. 2B**). These observations are similar to previous reports of spindle bending and breaking in fission yeast cells with chemically inhibited fatty acid synthesis (Takemoto et al., 2016). Our data are therefore compatible with the hypothesis that fatty acid synthesis mutants are prone to mitotic defects as a result of insufficient supply of membrane phospholipids needed for the nuclear envelope expansion during anaphase (Makarova et al., 2016; Takemoto et al., 2016). However, as we show below, other factors also contribute to the mitotic defects in *cbf11Δ* cells.

### Inactivation of the spindle assembly checkpoint does not affect the incidence of catastrophic mitosis in *cbf11Δ* cells

The increased duration of the pre-anaphase mitotic period in *cbf11Δ* cells (**Fig. 2B**) prompted us to investigate the role of the spindle assembly checkpoint (SAC) in mitotic fidelity of the *cbf11Δ* mutant. Through APC/C inactivation, SAC inhibits premature sister chromatid separation and mitotic exit until all chromosomes are properly attached to the spindle microtubules (Musacchio, 2015). Theoretically, SAC activation could provide *cbf11Δ* cells with more time to deal with any mitotic problems they might encounter. Under such a scenario, genetic perturbation of the SAC mechanism should lead to a higher incidence of unresolved mitotic defects. On the other hand, a failure to silence the SAC after chromosomes have been successfully attached to the spindle might result in prolonged metaphase arrest and perturbed timing of downstream mitotic and cytokinetic events. Therefore, under this alternative scenario, SAC abolition might actually suppress the mitotic defects of the *cbf11Δ* strain. However, when we inactivated SAC in the *cbf11Δ* background by deleting either *bub1* or *mad2*, we have not observed any significant changes in the incidence of catastrophic mitosis compared to *cbf11Δ* single mutants (**Fig. 3A, B**). These results suggest that SAC activity only plays a minor role (if any) in the mitotic defects observed in *cbf11Δ* cells.

**Figure 3.**
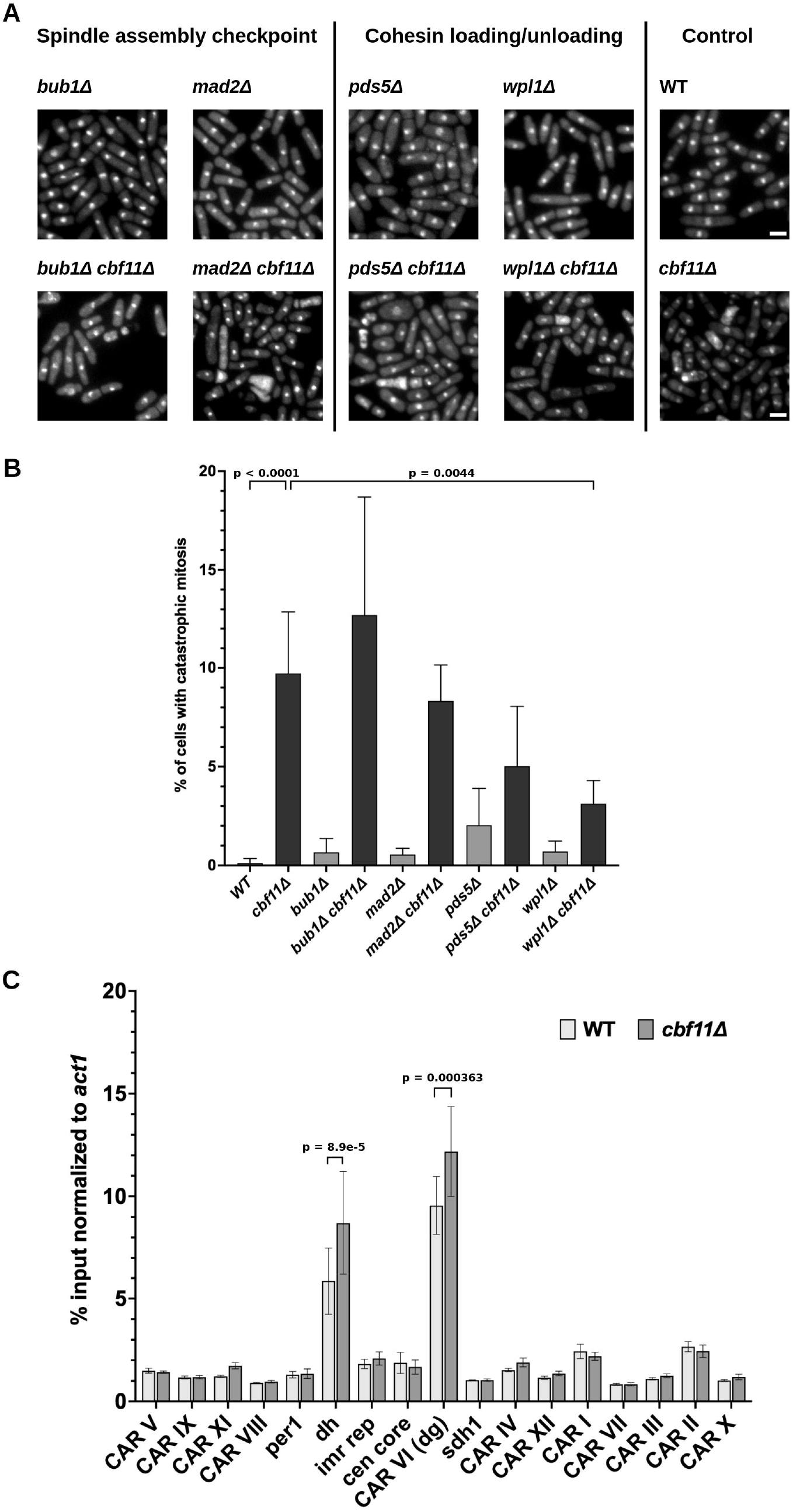
Suppression of catastrophic mitosis in *cbf11Δ* cells by mutation in cohesin loader. Representative microscopic images of cells of the indicated genotypes. Nuclei were stained with DAPI. Bar represents 5 μm. **(B)** Incidence of catastrophic mitosis in the indicated strains. At least 400 cells were scored per sample; values represent means + SD from 3 independent experiments. The significance of suppression of catastrophic mitosis in the respective double mutants compared to the *cbf11Δ* single mutant was determined by oneway ANOVA followed by the Bonferroni multiple comparison test; non-significant comparisons are not indicated for clarity. **(C)** Psm1-GFP occupancy at CARs was determined by ChIP-qPCR and normalized to the control *act1* locus. Values represent means + SD from 3 independent experiments. The significance of observed differences was determined by multiple unpaired t-tests followed by the Bonferroni post-test; non-significant comparisons are not indicated for clarity.

### Mitotic defects in *cbf11Δ* cells are partially suppressed by cohesin-loader mutations

The dynamics of sister chromatid cohesion are crucial for the successful execution of mitosis. Interestingly, the deletion of *cbf11* was found to have negative genetic interactions with mutations in the Mis4 cohesin loading factor (adherin; *mis4-242*) and the Eso1 cohesin N-acetyltransferase (*eso1-G799D*), which are both important for the establishment of cohesion (Chen et al., 2012). Therefore, we tested whether cohesin plays any role in the mitotic defects of *cbf11Δ* cells. We first quantified cohesin (Psm1) localization to the known major cohesin-associated regions (CARs) on chromosome II (Bhardwaj et al., 2016) in asynchronous WT and *cbf11Δ* cells using ChIP-qPCR (**Fig. 3C**). While cohesin occupancy along the chromosome arms was similar in both strains, we found significantly higher cohesin occupancy at the centromeric *dh* and *dg* repeats. Notably, centromeres are the regions where sister chromatin cohesion is abolished last during mitosis (Peters et al., 2008). Since *cbf11Δ* cells show altered cell-cycle and pre-anaphase mitotic duration compared to WT (**Fig. 2**), the observed difference in cohesin occupancy might merely reflect these changes in the timing of cell cycle progression. Alternatively, altered cohesin dynamics could play a role in the *cbf11Δ* mitotic defects. To test this hypothesis, we created double mutants of *cbf11Δ* and factors involved in cohesin un/loading (*pds5Δ, wpl1Δ*) (Murayama and Uhlmann, 2015; Tanaka et al., 2001) and determined their rates of catastrophic mitosis (**Fig. 3A, B**). Notably, the deletion of *wpl1* resulted in significant suppression of the mitotic defects compared to the *cbf11Δ* single mutant, and a noticeable but statistically not significant decrease was also observed upon deletion of *pds5*. It is, therefore, possible that altered timing of cohesin loading and/or removal and the ensuing changes in sister chromatid cohesion contribute to the incidence of catastrophic mitosis in cells lacking Cbf11.

We also used our panel of *cbf11Δ* double mutants to test whether suppression of mitotic defects correlated with suppression of slow growth (**Fig. S2**) and abnormal cell morphology (**Fig. 3A**). While we did not observe any improvements in growth rate or cell morphology in the *cbf11Δ* double mutants with *mad2Δ, bub1Δ* or *pds5Δ*, we found that the *cbf11Δ wpl1Δ* double mutant also showed suppression of aberrant cell morphology, but not of growth defects. Therefore, these three types of defects (mitotic defects, slow growth, abnormal cell morphology) of *cbf11Δ* cells are at least partially independent of each other and can be separated genetically.

### Centromeric chromatin is perturbed in cells lacking Cbf11

A previous high-throughput screen suggested that *cbf11Δ* cells are sensitive to the microtubule poison thiabendazole (TBZ) (Han et al., 2010). Furthermore, other studies showed that the loss of *cbf11* displays intriguing genetic interactions with mutations in chromatin- and kinetochore-related factors. Namely, *cbf11Δ* interacts negatively with the deletion of *sgf73*, a subunit of the SAGA histone acetyltransferase complex, which shows decreased histone acetylation at H3K9 and H3K16 (Guo et al., 2014). Also, *cbf11Δ* interacts negatively with the deletion of the *gsk3* kinase gene (Rallis et al., 2017). Notably, Gsk3 is important for localization of the Mis12 protein to the kinetochore and, subsequently, proper chromosome segregation during mitosis (Goshima, 2003). This prompted us to analyze the centromeric chromatin in *cbf11Δ* cells in more detail.

We first validated the previous report of *cbf11Δ* cells being sensitive to TBZ. When plated on YES medium containing TBZ, the *cbf11Δ* mutant indeed showed strong sensitivity to the drug (**Fig. S3A**), in line with the lowered robustness of spindle functioning in these cells (**Fig. 1B, Fig. S1, Fig. 2B-D**). The fission yeast centromeres consist of a central core, where histone H3 is replaced with the CENP-A histone variant, and inner and outer repeats that are heterochromatinized, hypoacetylated, and feature methylation of H3K9 (Kniola et al., 2001; Pidoux and Allshire, 2004). Importantly, perturbations of the centromeric heterochromatin lead to mitotic defects (Allshire et al., 1995; Roche et al., 2016). To assess the status of centromeric and pericentromeric chromatin in *cbf11Δ* cells, we performed ChIP-seq analysis of H3K9 acetylation (H3K9ac; target of SAGA) and dimethylation (H3K9me2). Compared to WT, the *cbf11Δ* mutant exerted an altered pattern of H3K9me2 occupancy with large centromeric regions being exceedingly hypermethylated. On the other hand, the H3K9ac pattern showed mild alterations in the pericentromeric regions and outer centromeric repeats (**Fig. 4A, Fig. S3B, C**). To test whether the altered distributions of histone marks had any impact on gene expression, we measured transcript levels of genes flanking centromere I (*per1, sdh1*) and of the centromeric *dh* and *dg* repeats (note that these repeats are present as multiple copies in all 3 fission yeast centromeres and individual repeats cannot be distinguished in our RT-qPCR assay). Indeed, we found that mRNA levels of *per1* and *sdh1* were ∼1.5x higher and ∼2x lower, respectively, in *cbf11Δ* cells compared to WT. Furthermore, the *dh* repeats showed significantly lower expression in the *cbf11Δ* mutant (**Fig. 4B**), possibly due to the hypermethylation of theses loci. Finally, we tested whether the fidelity of chromosome segregation during mitosis was negatively affected in *cbf11Δ* cells. To this end, we analyzed the rate of loss of a non-essential minichromosome (Ch16) derived from chromosome III (Tinline-Purvis et al., 2009). We found that *cbf11Δ* cells had a ∼9-fold higher rate of Ch16 loss per generation compared to WT (**Fig. 4C**). Collectively, these results suggest that the organization and function of (peri)centromeric chromatin is perturbed in cells lacking Cbf11 and that this might contribute to their problems with chromosome segregation.

**Figure 4.**
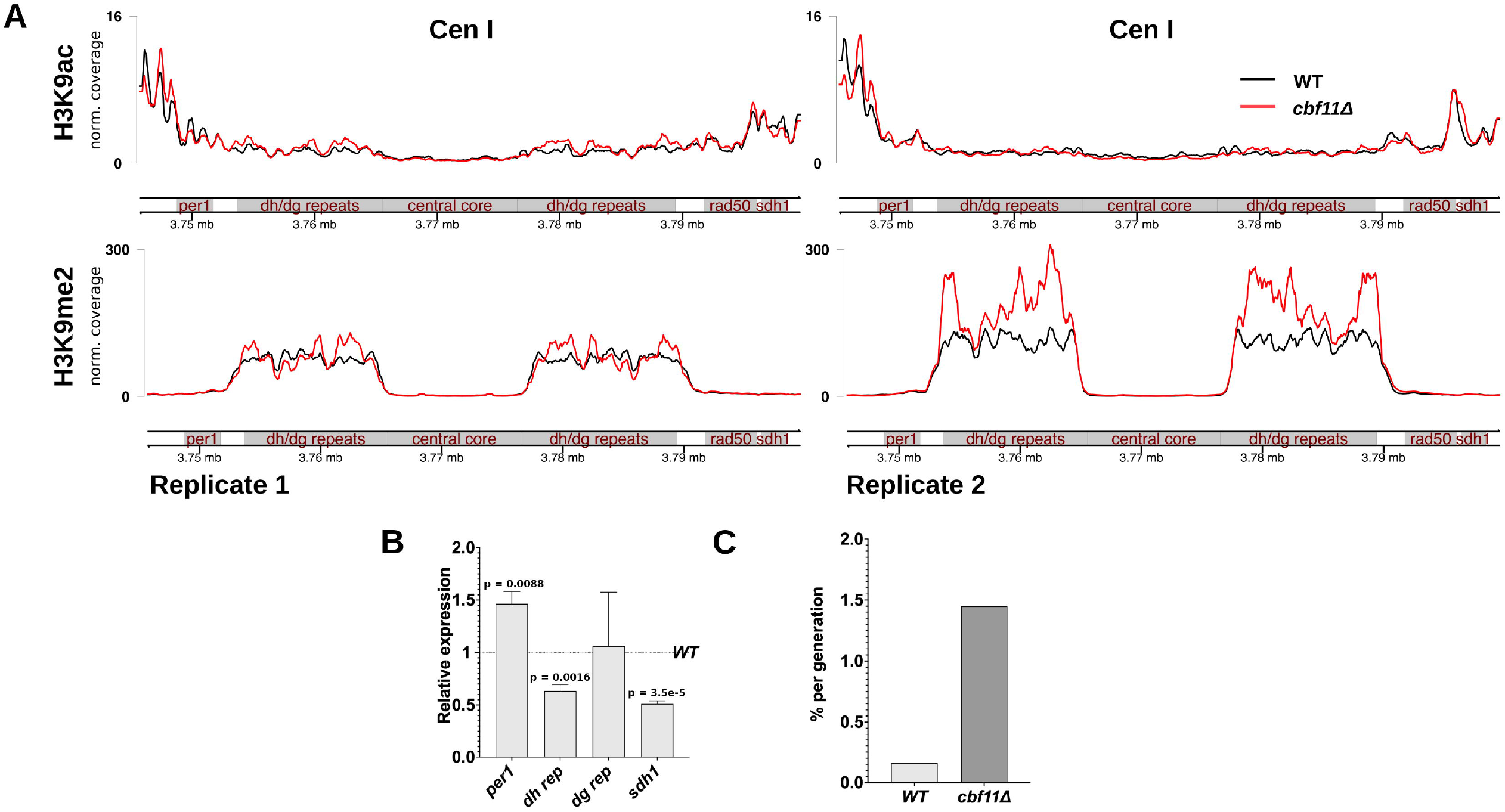
Cells lacking Cbf11 show perturbed centromeric chromatin. **(A)** Normalized ChIP-seq coverage of histone H3K9 acetylation and dimethylation at centromere I in WT and *cbf11Δ* cells. Data from 2 independent experiments are shown. **(B)** Expression of centromeric and pericentromeric genes at centromere I in *cbf11Δ* cells (normalized to expression in WT). Values represent means + SD from 3 independent experiments. The significance of observed differences was determined by multiple unpaired t-tests followed by the Bonferroni post-test; non-significant comparisons are not indicated for clarity. **(C)** Rate of Ch16 minichromosome loss per generation (3752 and 1856 colonies were scored for WT and *cbf11Δ* cells, respectively).

## DISCUSSION

The requirement for fatty acid synthesis for the closed mitosis of *S. pombe* was noted decades ago (Saitoh et al., 1996). But it has been only recently suggested that this requirement is connected to adequate membrane phospholipid supply for the rapid nuclear envelope expansion during the anaphase of closed mitosis (Makarova et al., 2016; Takemoto et al., 2016; Zach and Převorovský, 2018). We have previously shown that the Cbf11 transcription factor regulates lipid metabolism genes, and its absence leads to increased occurrence of catastrophic mitosis (“cut” phenotype) (Převorovský et al., 2009; Převorovský et al., 2016). However, since Cbf11 regulates other genes and processes apart from lipid metabolism (Převorovský et al., 2015), we set out to characterize in detail the mitotic defects of the *cbf11Δ* mutant in order to assess whether other factors might also contribute to the mitotic defects observed in this mutant. While our results are compatible with fatty acid shortage contributing to mitotic defects, we identified several additional factors, not directly related to lipid metabolism, that affect mitotic fidelity in *cbf11Δ* cells. Specifically, these factors are related to (peri)centromeric chromatin structure, gene expression, and sister chromatid cohesion dynamics.

First, we found that it takes the *cbf11Δ* cells longer to finish all pre-anaphase mitotic processes, suggesting that problems are present even before the insufficient supply of membrane phospholipids can take an effect on the closed mitosis (**Fig. 2A, B**). This indicates that proper microtubule attachment to kinetochores might be compromised and takes longer to achieve in *cbf11Δ* cells, possibly triggering the SAC. Intriguingly, SAC inactivation has been shown to suppress the temperature sensitivity of the *cut9-665* APC/C mutant, which is also prone to catastrophic mitosis (Elmore et al., 2014). However, SAC inactivation had very little effect on mitotic fidelity in the *cbf11Δ* mutant (**Fig. 3A, B**), so either the defects lie in other pre-anaphase step(s), or SAC is unable to sense and/or respond to kinetochore attachment defects in this particular case.

Second, the mitotic spindle dynamics is compromised in *cbf11Δ* cells, as cells undergoing aberrant mitosis showed lower spindle elongation rate (**Fig. 2C, Fig. S1B**), and spindle bending and/or breaking was also observed, together with chromosome detachment from the spindle during anaphase (**Fig. 1B, Fig. 2B**). Furthermore, *cbf11Δ* cells are sensitive to the microtubule poison TBZ (**Fig. S3A** and (Han et al., 2010)) and are prone to chromosome loss (**Fig. 4C**). These observations once again suggest that kinetochore function might be compromised when Cbf11 is missing. Nevertheless, it is also conceivable that the slower spindle elongation and spindle bending can be caused by the physical forces exerted by the nuclear envelope that expands too slowly, out of sync with the growing spindle (Takemoto et al., 2016).

Third, the deletion of *cbf11* was shown to exert negative genetic interactions with mutations in two essential factors involved in cohesin loading, Eso1 (*eso1-G799D*) and Mis4 (*mis4-242*) (Chen et al., 2012), suggesting that cohesin loading and sister chromatid cohesion might be compromised in *cbf11Δ* cells. However, we did not detect any decrease in cohesin occupancy along chromosome arms, and we actually found increased cohesin occupancy at the centromeric *dh*/*dg* repeats in this mutant (**Fig. 3C**). Moreover, deletion of the cohesin un/loader Wpl1/WAPL partially suppressed the mitotic defects of *cbf11Δ* cells (**Fig. 3B**). Notably, deletion of *wpl1* can also rescue the otherwise lethal deletion of *eso1* (Feytout et al., 2011), suggesting the requirement for a fine balance between cohesin loading and removal from DNA during the cell cycle. Collectively, these results indicate that altered cohesin dynamics, in general, contribute to the mitotic defects of *cbf11Δ* cells.

Finally, the above-mentioned centromere-related defects of *cbf11Δ* cells are accompanied by altered chromatin marks at and around the centromeres (**Fig. 4A, Fig. S3B, C**), together with altered gene expression from these regions (**Fig. 4B**). Specifically, H3K9 tends to be exceedingly hypermethylated at the *dh*/*dg* centromeric repeats (which also show increased cohesin occupancy), and these repeats have lower expression in *cbf11Δ* cells compared to WT. Importantly, a mutation in the SAGA histone acetyltransferase complex, which leads to decreased H3K9 acetylation (Nugent et al., 2010) and thus presumably increased H3K9 methylation (Alper et al., 2013; Nakayama et al., 2001), shows a negative genetic interaction with *cbf11Δ*. This suggests that such a combination of two hypermethylation-promoting mutations is deleterious for the cells, highlighting chromatin structure as yet another important contributor to the low mitotic fidelity of *cbf11Δ* cells. While centromeric heterochromatin (H3K9me2) is required for proper chromosome segregation (Ekwall et al., 1996), heterochromatin is a dynamic structure that undergoes tightly regulated re-establishment in every cell cycle (Chen et al., 2008), and it is conceivable that exceedingly high levels of H3K9 methylation, as observed in *cbf11Δ* cells, might disrupt the cyclic behavior of centromeric chromatin structure and function. It is also important to note that heterochromatin, kinetochore function, cohesin occupancy, and gene expression are all interconnected and actually interdependent (Bernard et al., 2001; Folco et al., 2019; Grewal and Jia, 2007; Gullerova and Proudfoot, 2008; Nonaka et al., 2002; Volpe et al., 2002). Therefore, our findings implicating all these structures and processes in the mitotic defects of *cbf11Δ* cells might potentially represent different facets of a common underlying root cause.

Another important question is whether the chromatin-related cause(s) of mitotic defects are specific to the *cbf11Δ* mutant, or they apply more generally to other mutants with perturbed lipid metabolism (Zach and Převorovský, 2018). While proper (centromeric) chromatin structure and dynamics might simply become more critical in cells compromised by insufficient membrane phospholipid supply, other possible explanations recently emerged. Fatty acid synthesis is a major consumer of acetyl-CoA, which also serves as a substrate for histone acetylation. Remarkably, metabolic manipulations leading to altered acetyl-CoA levels have been shown to trigger changes in histone modifications and expression of specific genes in the budding yeast and mammals (Galdieri and Vancura, 2012; McDonnell et al., 2016; Takahashi et al., 2006; Wellen et al., 2009). Moreover, specific changes in chromatin structure and gene expression can be brought about by highly localized alterations in acetyl-CoA availability in the nucleus (Mews et al., 2017). It is thus possible that in lipid metabolism mutants the chromatin landscape is altered in such a way that, for example, a subset of genes becomes overexpressed (as in the case of *per1* in *cbf11Δ* cells; **Fig. 4B**) while centromeric heterochromatin is exceedingly hypermethylated, and this imbalance leads to defective mitosis. In summary, while the exact mechanism of how lipid metabolism is linked to centromeric chromatin function and fidelity of closed mitosis needs to be addressed by future studies, we have demonstrated several novel factors, not directly related to lipid metabolism, that affect mitotic fidelity in cells with perturbed lipid homeostasis. Moreover, since closed mitosis is typical for many clinically and economically important fungal species, our findings could inform research in the fields of drug design and pest control.

## MATERIALS AND METHODS

### Cultivation media and construction of strains

Standard methods and media were used for the cultivation of *Schizosaccharomyces pombe* strains (Sabatinos and Forsburg, 2010). Unless indicated otherwise, *S. pombe* strains were grown at 32°C in YES with the appropriate supplements or stressors, as required.

The strains used in the study are listed in **Table S1**. Deletion of *cbf11* was carried out using the pMP91 targeting plasmid based on pCloneNAT1 as described (Gregan et al., 2006). All other strains used in this study were constructed by standard genetic crosses.

### Growth rate measurements

*S. pombe* cells were pre-cultured to exponential phase in 180 μl of YES in 96-well culture plates at 32°C in the VarioSkan Flash plate reader (Thermo Scientific). 5 μl of the preculture were used to inoculate 175 μl of fresh YES in 96-well culture plates and incubated for ∼48 hours in the VarioSkan Flash plate reader at 32°C with background shaking (1080 spm, rotation diameter 1 mm, a measuring time of 100 ms and 5 nm bandwidth). Optical densities were measured at 10 min intervals at λ = 600 nm. Culture doubling times were calculated in R using the “growthrates” package (https://github.com/tpetzoldt/growthrates).

### Chromosome loss assay

WT and *cbf11Δ* cells harboring the Ch16-RMGAH non-essential minichromosome (Hylton et al., 2020; Tinline-Purvis et al., 2009) were grown on YES plates with low adenine content (10 mg/l). Under such conditions, cells that have lost the minichromosome and thus are adenine auxotrophs (*ade6-M210*) turn dark red. Sectored colonies (at least half-red) lost the minichromosome in the very first cell division after plating and are used to calculate the per-division chromosome loss rate. Fully red colonies lost the minichromosome prior to plating and are excluded from the analysis (Javerzat, 1996).

### ChIP-qPCR

For chromatin immunoprecipitation (ChIP), 200 ml of fission yeast cultures were grown in YES to the density of 0.5 - 1.2×10^7^ cells/ml, fixed with 1% formaldehyde for 30 min, quenched with 10 ml of 2.5 M glycine for 10 min, washed with water and broken with glass beads in Lysis Buffer (50 mM HEPES, 1 mM EDTA, 150 mM NaCl, 1% Triton X-100, 0.1% sodium deoxycholate, FY protease inhibitors (Serva); pH 7.6) using the FastPrep-24 instrument (MPI Biomedicals). Chromatin extracts were sheared with the Bioruptor sonicator (Diagenode) to yield DNA fragments of ∼200 bp; 1/10 of the total chromatin extract amount was kept for input DNA control. 5 μg of anti-GFP antibody (ab290, Abcam) was added to ∼1 mg of chromatin extract and incubated for 1 hour at 4°C. Antibody-chromatin complexes were then captured with Dynabeads® Protein A (cat. no. 10002D, Life technologies). The precipitated material was washed twice with Lysis Buffer, Lysis 500 Buffer (50 mM HEPES, 1 mM EDTA, 500 mM NaCl, 1% Triton X-100, 0.1% sodium deoxycholate; pH 7.6), LiCl/NP-40 Buffer (10 mM Tris-HCl, 1 mM EDTA, 250 mM LiCl, 1% Nonidet P-40, 1% sodium deoxycholate; pH 8.0), once in TE (10 mM Tris-HCl, 1 mM EDTA; pH 8.0), and eluted in Elution Buffer (50 mM Tris-HCl, 10 mM EDTA, 1% SDS; pH 8.0). Crosslinks were reversed overnight at 65°C, samples were treated with DNase-free RNase followed by proteinase K, and purified twice using phenol-chloroform extraction and sodium acetate precipitation. The enrichment of specific target DNA sequences and controls in the immunoprecipitated material vs the input was determined by quantitative PCR (qPCR) using the HOT FIREPol® EvaGreen® qPCR Supermix (Solis BioDyne) and the LightCycler 480 II instrument (Roche). The primers used are listed in **Table S2**.

### ChIP-seq

ChIP was performed analogously as in ChIP-qPCR with the following modifications: Harvested cells were washed in PBS. For each immunoprecipitation, 5 μg of antibody (H3K9ac: Ab4441, H3K9me2: Ab1220, all Abcam) were incubated with the chromatin extract for 1 hour at 4°C with rotation. Immunoprecipitated and input DNA samples were purified using phenol-chloroform extraction and sodium acetate/ethanol precipitation. In biological replicate 2, DNA purification on AMPure XP beads (Beckman Coulter, AC63880) was performed following the phenol-chloroform extraction to remove low-molecular fragments and RNA. Library construction and sequencing were performed by BGI Tech Solutions (Hong Kong) using the BGISEQ-500 sequencing system (∼24 M of 50 nt single end reads per sample).

The *S. pombe* reference genome sequence and annotation were obtained from PomBase (release date 2018-09-04) (Lock et al., 2019; Wood et al., 2002). Read quality was checked using FastQC version 0.11.8 (https://www.bioinformatics.babraham.ac.uk/projects/fastqc/), and reads were aligned to the *S. pombe* genome using HISAT2 2.1.0 (Kim et al., 2019) and SAMtools 1.9 (Bonfield et al., 2021; Li et al., 2009). Read coverage tracks (i.e., target protein occupancy) were then computed and normalized to the respective mapped library sizes using deepTools 3.3.1 (Ramírez et al., 2016). Plots of coverage tracks were made in R using the Gviz package (Hahne and Ivanek, 2016). The raw ChIP-seq data are available from the ArrayExpress (https://www.ebi.ac.uk/arrayexpress/) database under the accession number E-MTAB-11081. The scripts used for ChIP-seq data processing and analysis are available from https://github.com/mprevorovsky/ox-stress_histones.

### RT-qPCR

Total RNA was extracted from exponentially growing *S. pombe* cells using the MasterPure Yeast RNA Purification Kit (Epicentre). The procedure for removal of DNA contamination was as per the specification of the manufacturer but performed twice to efficiently remove the DNA from the samples. Purified RNA was converted to cDNA using random primers and the RevertAid™ Reverse Transcriptase kit (Fermentas) using the protocol specified by the manufacturer. Quantitative PCR was performed using the HOT FIREPol® EvaGreen® qPCR Supermix (Solis BioDyne) and the LightCycler 480 II instrument (Roche). For RT-qPCR, *act1* (actin) and/or *rho1* (Rho GTPase) were used as reference genes. The primers used are listed in **Table S2**.

### Microscopy and image analysis

For nuclear staining, exponentially growing *S. pombe* cells were pelleted by centrifugation (1000 x g, 3 min, 25°C) and fixed in 70% ethanol. Fixed cells were centrifuged again and rehydrated in deionized H_2_O. Cells were stained with 0.5 μg/ml 4’,6-diamidine-2’-phenylindole dihydrochloride (DAPI). Images were taken using the Olympus CellR microscopic system with a 60x objective. The frequency of catastrophic mitosis occurrence was determined by manual evaluation of microscopic images using ImageJ software, version 1.52p (Schneider et al., 2012). At least 400 cells were analyzed per sample.

For visualizing live-cells, exponentially growing *S. pombe* cells were collected by centrifugation (1000 x g, 3 min, 25°C) and resuspended in ∼5 μl of YES. 1 μl of cell suspension was applied on 2% agarose-YES media solidified on a PDMS spacer (generously provided by Phong Tran, Institut Curie, Paris) and covered with a coverslip. Cells were imaged using the Olympus CellR microscope with automatic z-axis objective movement with a 60x objective fitted with a 12-bit monochromatic Hamamatsu ORCA CCD camera. The images captured consisted of 10**-**11 Z-stacks of 0.3**-**0.5 μm sections, 500 ms exposure in red and green channels with a 2 min cycle, and 2×2 binning. Images were then processed using ImageJ 1.52p (Schneider et al., 2012) with maximum intensity projection. A custom ImageJ macro script was used for automating the image processing. Nuclear and spindle dynamics were analyzed using the tools available in ImageJ. The nuclear distance was measured by using Hht2–GFP signals and converting the green channel images to binary, plotting the profile, and measuring the maximum section. Spindle length was calculated using mCherry-Atb2 signals. Spindle elongation rate was calculated by computing the slope from the exponential region of the spindle length vs time (see **Supplementary Figure 1**). Statistical analyses and generating of figures were performed in R studio and Graphpad Prism.

## ACKNOWLEDGEMENTS

This work was supported by the Charles University grant no. PRIMUS/MED/26. We are very grateful to Phong Tran for his advice and material support for live-cell microscopy. We thank Karl Ekwal, Charalampos Rallis, Tim Humphrey, and Phong Tran for providing strains. The *bub1Δ, mad2Δ, pds5Δ, wpl1Δ*, and *psm1-GFP* strains were provided by The Yeast Genetic Resource Center Japan. We also thank Adéla Kracíková for technical assistance and Ondřej Šebesta for assistance with microscopy. Microscopy was performed in the Laboratory of Confocal and Fluorescence Microscopy co-financed by the European Regional Development Fund and the state budget of the Czech Republic, projects no. CZ.1.05/4.1.00/16.0347 and CZ.2.16/3.1.00/21515, and supported by the Czech-BioImaging large RI project LM2018129. Computational resources were supplied by the project “e-Infrastruktura CZ” (e-INFRA LM2018140) provided within the program Projects of Large Research, Development, and Innovations Infrastructures.

