## Supplemental Data for "Altered cohesin dynamics and histone H3K9 modifications contribute to mitotic defects in the *cbf11Δ* lipid metabolism mutant"

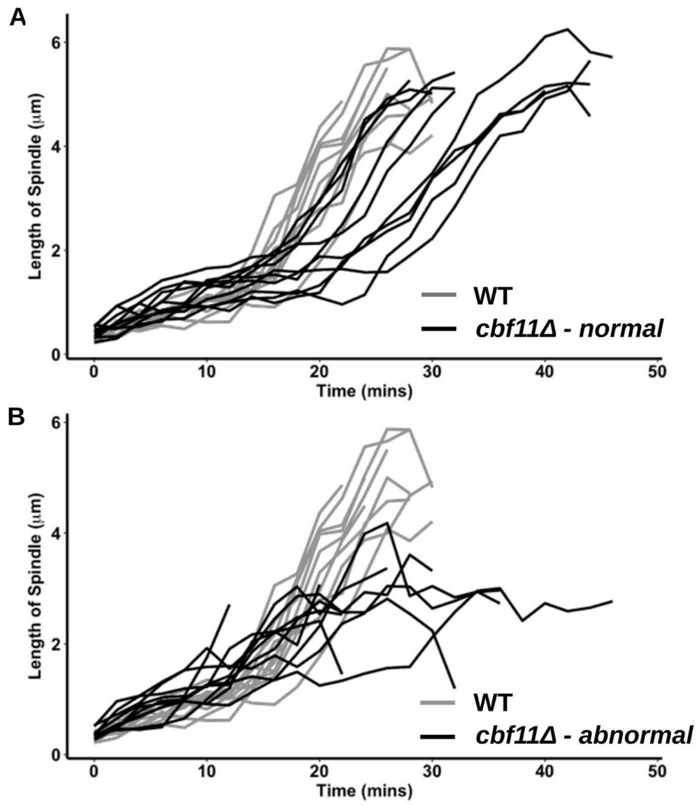


**Supplementary Figure 1 – Mitotic spindle dynamics in WT and *cbf11Δ* cells.**
**(A, B)** Spindle length in live mitotic cells of the WT and *cbf11Δ* strains was monitored over time. The time scale corresponds to **Fig. 2A, B**, starting at the “anaphase” mark. Ten cells were analyzed for each indicated group.

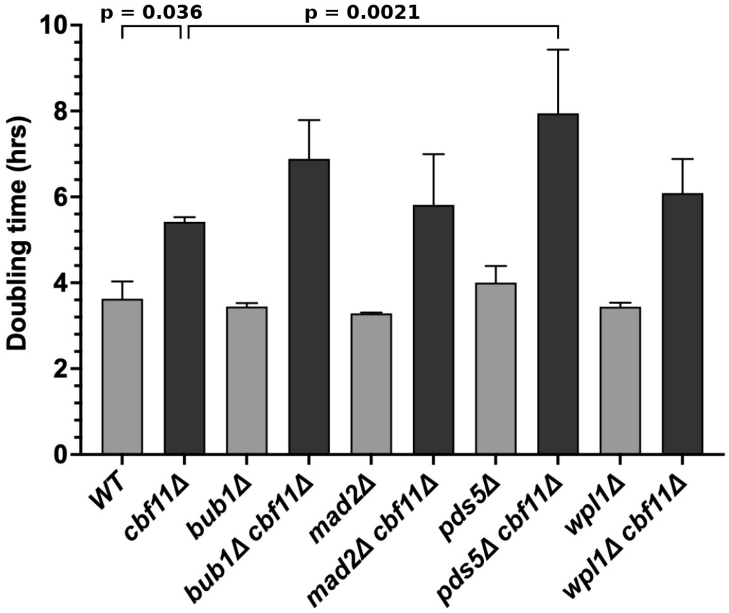


**Supplementary Figure 2 - Doubling times of *cbf11Δ* double mutants.**
Doubling times of cultures growing exponentially in YES medium at 32°C. Values represent means + SD from 3 independent experiments. The significance of differences versus the *cbf11Δ* single mutant was determined by one-way ANOVA followed by the Bonferroni multiple comparison test; non-significant comparisons are not indicated for clarity.

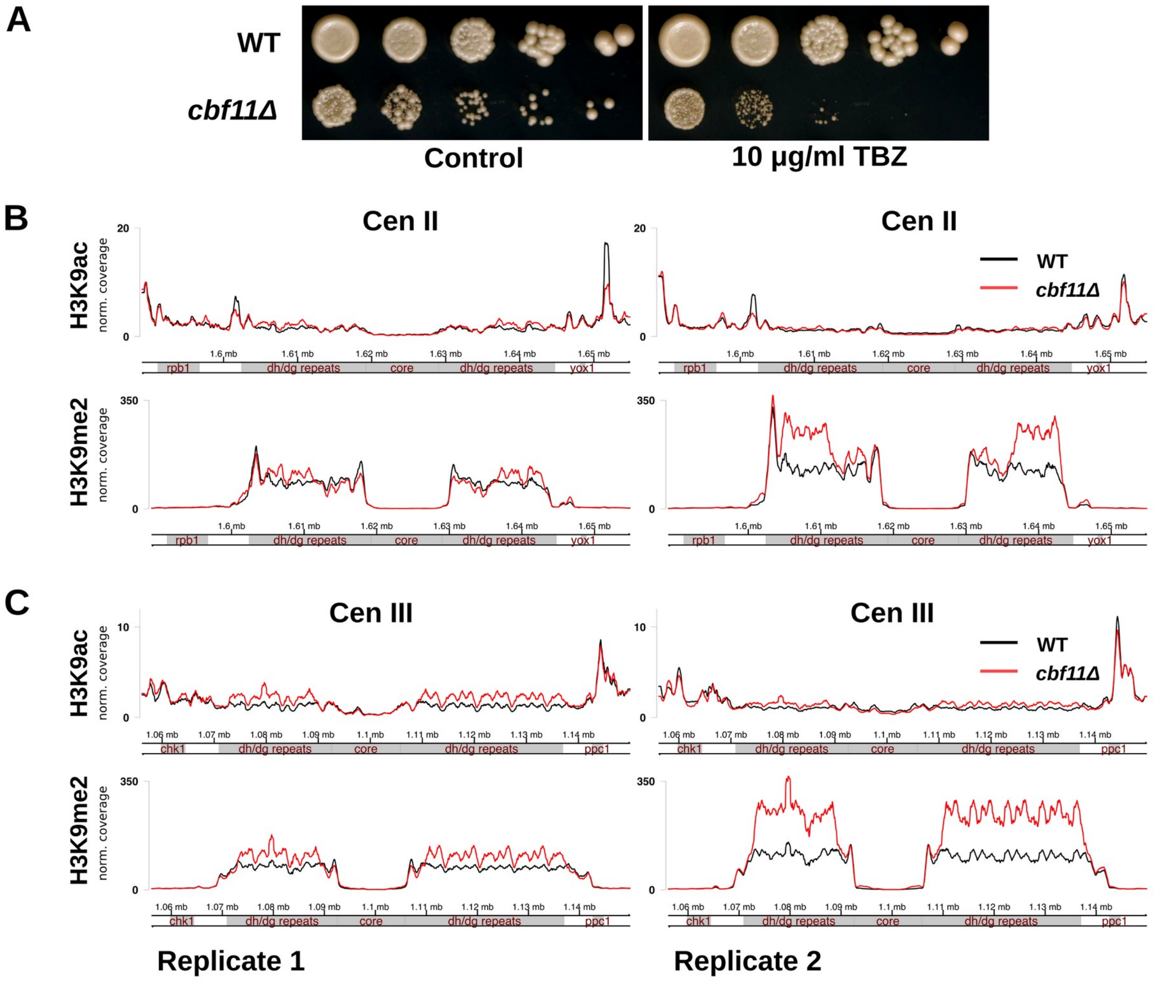


**Supplementary Figure 3 – Perturbed centromeric chromatin in *cbf11Δ* cells.**

**(A)** Tenfold serial dilutions of cells were spotted on the indicated plates and incubated at 32°C. **(B, C)** Normalized ChIP-seq coverage of H3K9 acetylation and dimethylation at centromeres II and III, respectively, in WT and *cbf11Δ* cells. Data from 2 independent experiments are shown.

**Supplementary Table 1 - List of strains used in this study.**

| Strain ID | Genotype | Source |
|---|---|---|
| JB22 | *972 wt h-* | Lab stock |
| JB32 | *h+s* | Lab stock |
| MP44 | *cbf11Δ::kanR, h+* | Lab stock |
| JB710 | *h- ura4-D18* | Lab stock |
| MP726 | *h- bub1Δ::KanR leu1* | YGRC NBRP, FY18581 |
| MP784 | *h- bub1Δ::KanR cbf11Δ::NatR leu1* | This study |
| MP730 | *h- pds5Δ::ura4 leu1 ura4D18* | YGRC NBRP, FY31429 |
| MP787 | *h- pds5Δ::ura4 cbf11Δ::NatR leu1 ura4D18* | This study |
| MP731 | *h- wpl1Δ::Hyg leu1* | YGRC NBRP, FY31430 |
| MP789 | *h- wpl1Δ::Hyg cbf11Δ::NatR leu1* | This study |
| MP732 | *h- mad2Δ leu1* | YGRC NBRP, FY9228 |
| MP790 | *h- mad2Δ cbf11Δ::NatR leu1* | This study |
| MP322 | *h- leu1 psm1-GFP:LEU2* | YGRC NBRP, FY11280 |
| MP745 | *h- leu1 psm1-GFP:LEU2 cbf11Δ::NatR* | This study |
| MP752 | *h+ mCherry-atb2-HygR hht2-GFP:Ura4+ ade6-21? leu1-32 ura4-D18* | Phong Tran |
| MP758 | *h+ mCherry-atb2-HygR hht2-GFP:Ura4+ cbf11Δ::NatR ade6-21? leu1-32 ura4-D18* | This study |
| JB814 | *h+ ura4-D18 leu1-32 ade6-M210 Ch16-RMGAH arg3-D4 his3-D1* | (Tinline-Purvis et al., 2009) |
| MP194 | *h+ ura4-D18 leu1-32 ade6-M210 Ch16-RMGAH Δcbf11::natR his3? Arg3?* | This study |

**Supplementary Table 2 - Primers used in this study.**

| ID | Sequence | Target, orientation | Source |
|---|---|---|---|
| RZ03 | TGGGTTCTGGTAGTTCAC | pre-centromeric region/*per1*, F | Lab stock |
| RZ04 | ACAGCAAGTTCACCTAAAG | pre-centromeric region/*per1*, R | Lab stock |
| RZ07 | CAAAAACTAAACGCCCATTC | *dh* repeats, F | Lab stock |
| RZ08 | CCGTGTTCTTTCCTTCTTC | *dh* repeats, R | Lab stock |
| MP180 | AATTGTGGTGGTGTGGTAATAC | *dg* repeats, F | Lab stock |
| MP181 | GGGTTCATCGTTTCCATTCAG | *dg* repeats, R | Lab stock |
| RZ13 | GAATGAGTGAGTGAGTACAAG | *imr* repeats, F | Lab stock |
| RZ14 | GCCTACAATTATACCCAATGAG | *imr* repeats, R | Lab stock |
| RZ27 | ATACGGAAAACTGAGACATC | Post-centromeric region/*sdh1, F* | Lab stock |
| RZ28 | GGACCAAATTCGCACTTC | Post-centromeric region/ sdh1, R | Lab stock |
| MP137 | TCCTCATGCTATCATGCGTCTT | *act1*, F | Lab stock |
| MP138 | CCACGCTCCATGAGAATCTTC | *act1*, R | Lab stock |
| MP169 | GGTCTATGTTCCCACTGTTT | *rho1*, F | Lab stock |
| MP170 | CTTCTTGTCCAGCCGTA | *rho1*, R | Lab stock |
| AV01 | AACCTCATCGAGAAGGAAA | Left-Centromere core, F | This study |
| AV02 | GAACTCCAGAAGGTGAGAA | Left-Centromere core, R | This study |
| AV03 | ACATAGCCTCTTACGTCTTT | Right-Centromere core, F | This study |
| AV04 | GAGTCGAACTTAGCAAATGG | Right-Centromere core, R | This study |
| AV05 | AAGGTATTAGTGGTCGGTTT | Centromere core central, F | This study |
| AV06 | CGCTTTCAGTGGTTGTTG | Centromere core central, R | This study |
| AV09 | GCTAGACTGATGATTCGCCCTTATA | Cohesin Associated Region I, F | This study; (Bhardwaj et al., |

| | | | |
|---|---|---|---|
| | | | 2016) |
| AV10 | TTTACCGGCGAGTTAGCAGAGTA | Cohesin Associated Region I, R | This study; (Bhardwaj et al., 2016) |
| AV11 | ATTCGTGGCAGATTTTGCTCT | Cohesin Associated Region II, F | This study; (Bhardwaj et al., 2016) |
| AV12 | AAACAAGCGTTTGGCAGTAGC | Cohesin Associated Region II, R | This study; (Bhardwaj et al., 2016) |
| AV13 | CTACTTCTGTTGCCTAATAGTAAAAACCGT | Cohesin Associated Region III, F | This study; (Bhardwaj et al., 2016) |
| AV14 | ACTGTCCGTGAATCAATCGTG | Cohesin Associated Region III, R | This study; (Bhardwaj et al., 2016) |
| AV15 | CAGGACAATTGGATGCTAAGTGTATT | Cohesin Associated Region IV, F | This study; (Bhardwaj et al., 2016) |
| AV16 | GGAACCTCTTCATTGCTTAAGCAT | Cohesin Associated Region IV, R | This study; (Bhardwaj et al., 2016) |
| AV17 | TTACTGAGGTTGGAAAGAAGGCAT | Cohesin Associated Region V, F | This study; (Bhardwaj et al., 2016) |
| AV18 | GGGTCTGCTTATAAAATTCGTGAAC | Cohesin Associated Region | This study; |

| | | | |
|---|---|---|---|
| | | V, R | (Bhardwaj et al., 2016) |
| AV19 | AAGACTGTTGTTGAGTGCTGTGGA | Cohesin Associated Region VI, F | This study; (Bhardwaj et al., 2016) |
| AV20 | CCATGCTTTTAGTGCGGTCA | Cohesin Associated Region VI, R | This study; (Bhardwaj et al., 2016) |
| AV21 | AGGCAGGAATCAGAAGGAGAGTT | Cohesin Associated Region VII, F | This study; (Bhardwaj et al., 2016) |
| AV22 | CAATCTCACCCACTTCTCCACTT | Cohesin Associated Region VII, R | This study; (Bhardwaj et al., 2016) |
| AV23 | GGAAGCCGATATCGGGTTAATATT | Cohesin Associated Region VIII, F | This study; (Bhardwaj et al., 2016) |
| AV24 | CCATGACAGTTCAGAAACCTCAT | Cohesin Associated Region VIII, R | This study; (Bhardwaj et al., 2016) |
| AV25 | GTTGCTTGCTTAACACTCACGAGTA | Cohesin Associated Region IX, F | This study; (Bhardwaj et al., 2016) |
| AV26 | CTGGTTCAATATCGATAATATACAAGCAG T | Cohesin Associated Region IX, R | This study; (Bhardwaj et al., 2016) |

| | | | |
|---|---|---|---|
| AV27 | GACACTTACAATATGGTGCTCTACGA | Cohesin Associated Region X, F | This study; (Bhardwaj et al., 2016) |
| AV28 | TGCGGTCAATTGTTTTCATCTAC | Cohesin Associated Region X, R | This study; (Bhardwaj et al., 2016) |
| AV29 | GGATACGTTCTATATGGTATGCATACGA | Cohesin Associated Region XI, F | This study; (Bhardwaj et al., 2016) |
| AV30 | ATCTTCACTCGCTTTTAAAGCCTGT | Cohesin Associated Region XI, R | This study; (Bhardwaj et al., 2016) |
| AV31 | AACCACCTATTCAAAGAGAGCTGCT | Cohesin Associated Region XII, F | This study; (Bhardwaj et al., 2016) |
| AV32 | TGTGTTTGCAGTATAAGATCTCTCCTC | Cohesin Associated Region XII, R | This study; (Bhardwaj et al., 2016) |
